# Alpha 1 Antitrypsin is an Inhibitor of the SARS-CoV-2–Priming Protease TMPRSS2

**DOI:** 10.1101/2020.05.04.077826

**Authors:** Nurit P. Azouz, Andrea M. Klingler, Victoria Callahan, Ivan V. Akhrymuk, Katarina Elez, Lluís Raich, Brandon M. Henry, Justin L. Benoit, Stefanie W. Benoit, Frank Noé, Kylene Kehn-Hall, Marc E. Rothenberg

## Abstract

Host proteases have been suggested to be crucial for dissemination of MERS, SARS-CoV, and SARS-CoV-2 coronaviruses, but the relative contribution of membrane versus intracellular proteases remains controversial. Transmembrane serine protease 2 (TMPRSS2) is regarded as one of the main proteases implicated in the coronavirus S protein priming, an important step for binding of the S protein to the angiotensin-converting enzyme 2 (ACE2) receptor before cell entry. The main cellular location where the SARS-CoV-2 S protein priming occurs remains debatable, therefore hampering the development of targeted treatments. Herein, we identified the human extracellular serine protease inhibitor (serpin) alpha 1 antitrypsin (A1AT) as a novel TMPRSS2 inhibitor. Structural modeling revealed that A1AT docked to an extracellular domain of TMPRSS2 in a conformation that is suitable for catalysis, resembling similar serine protease–inhibitor complexes. Inhibitory activity of A1AT was established in a SARS-CoV-2 viral load system. Notably, plasma A1AT levels were associated with COVID-19 disease severity. Our data support the key role of extracellular serine proteases in SARS-CoV-2 infections and indicate that treatment with serpins, particularly the FDA-approved drug A1AT, may be effective in limiting SARS-CoV-2 dissemination by affecting the surface of the host cells.

**Summary:** Delivery of extracellular serine protease inhibitors (serpins) such as A1AT has the capacity to reduce SARS-CoV-2 dissemination by binding and inhibiting extracellular proteases on the host cells, thus, inhibiting the first step in SARS-CoV-2 cell cycle (i.e. cell entry).

## Introduction

The COVID-19 pandemic is caused by the severe acute respiratory syndrome (SARS) - coronavirus (CoV) 2. The efficient transmission of this virus has led to exponential growth in the number of worldwide cases. Similar to other coronaviruses, SARS-CoV-2 entry into host cells relies on the proteolytic processing of spike (S) protein by host proteases and engagement of the angiotensin-converting enzyme 2 (ACE2) receptor (Hoffmann et al., 2020). Several proteases are crucial to coronavirus viral entry, and they are found at different subcellular locations. The S protein cleavage may occur extracellularly near the plasma membrane by cell surface proteases or intracellularly by lysosomal endopeptidase enzymes, such as cathepsin L, that facilitate viral entry by activating membrane fusion and subsequent cell entry through endocytosis, as in the case of MERS-CoV (Qing et al., 2020). Despite intensive research of the SARS-CoV-2 life cycle, the cellular location where the SARS-CoV-2 S protein priming occurs remains debatable. In particular, the interplay between the extracellular and intracellular proteases in the membrane fusion and cell entry of SARS-CoV-2 is controversial.

Transmembrane serine protease 2 (TMPRSS2), a cell surface serine protease, may be involved in cell entry of SARS-CoV-2. TMPRSS2 has been shown to cleave ACE2 at arginine and lysine residues within ACE2 amino acids 697–716, which enhances cell entry (Heurich et al., 2014). TMPRSS2 increases the entry of another coronavirus SARS-CoV, not only by processing of the S protein, but also by processing of the host receptor ACE2 (Heurich et al., 2014). Consistent with these findings, TMPRSS2-deficient mice have decreased viral spread of SARS-CoV in the airways compared to that of control mice (Iwata-Yoshikawa et al., 2019). In addition, the drug camostat mesylate (camostat), which inhibits a number of proteases including TMPRSS2, was shown to inhibit SARS-CoV-2 entry into cells in vitro. (Hoffmann et al., 2020; Matsuyama et al., 2020). Therefore, TMPRSS2 is regarded as one of the most important proteases for S protein priming and cell entry of SARS-CoV-2.

Herein, we developed a cell-based assay to identify TMPRSS2 inhibitors. We compared the efficiency of TMPRSS2 inhibition by synthetic and natural serine protease inhibitors that are cell permeable or have extracellular function, including drugs with known function as protease inhibitors. We identified alpha 1 antitrypsin (A1AT) as a novel inhibitor of TMPRSS2. Structural modelling of the Michaelis complex between TMPRSS2 and A1AT indicated that they dock on the cell surface in a conformation that is suitable for catalysis, resembling similar serine protease–inhibitor complexes. We further provided proof of concept for the potential utility of A1AT as an antiviral agent in SARS-CoV-2 infection. A1AT decreased SARS-CoV-2 copy number within target cells when applied during infection. The effect of A1AT was comparable to the effect of camostat, which was previously shown to inhibit SARS-CoV-2 cell entry (Hoffmann et al., 2020). In contrast to camostat, which is a cell permeable drug (Reihill et al., 2016), A1AT is a circulating extracellular protein that inhibits extracellular proteases and does not possess access to intracellular proteases (Strnad et al., 2020). These findings emphasize the importance of extracellular proteases to viral cell entry. We suggest that by inhibiting extracellular proteolytic activity, A1AT can potentially inhibit S protein processing and limit SARS-CoV-2 cell-cell spread and dissemination.

## Results

### Overexpressing TMPRSS2 and measuring proteolytic activity

We aimed to establish an experimental framework for quantifying TMPRSS2 proteolytic activity. We chose to overexpress TMPRSS2 in a human cell line, HEK-293T, because of its high transfectability. Western blot analysis of the cell lysates revealed a band at ∼60 kD in TMPRSS2-transfected cells but not in control cells (Figure 1A). GAPDH was used as a loading control. Measurements of the proteolytic activity of the transfected cells using the fluorogenic peptide substrate Boc-Gln-Ala-Arg-7-Amino-4-methylcoumarin (BOC-QAR-AMC) revealed a >2.5-fold increase in the proteolytic activity of the TMPRSS2-transfected cells compared with that of control cells (p = 0.0002; Figure 1B). The proteolytic activity of the TMPRSS2-transfected cells increased over time compared with that of control cells (Figure 1C). The mean proteolytic rate per minute of the TMPRSS2-transfected cells was increased by >3.5 fold compared to the proteolytic rate of control cells (p < 0.0001, Figure 1D). The proteolytic activity could be measured hours after addition of the substrate, and the average proteolytic activity per hour was increased by >3 fold in TMPRSS2-overexpressing cells compared with control cells (p < 0.0001, Figure 1E). Using serial dilutions of recombinant TMPRSS2, we estimated that the amount of TMPRSS2 that is expressed by TMPRSS2-overexpressing cells is about 100 ng/well (Supplementary Figure 1). These collective data demonstrated that overexpression of TMPRSS2 resulted in overproduction of functional TMPRSS2 and established an experimental system for accurately measuring the proteolytic activity of TMPRSS2.

**Figure 1.**
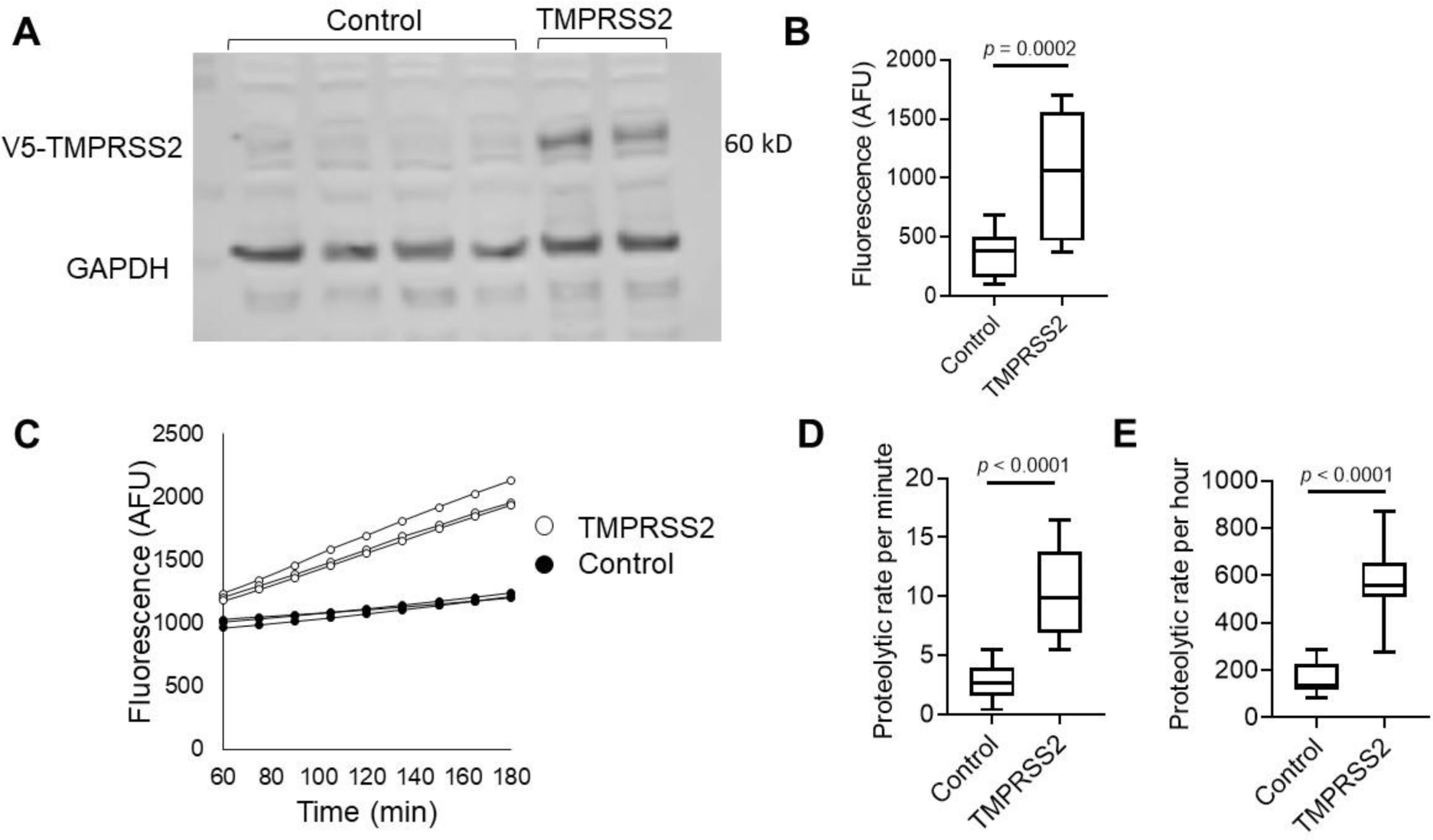
Measurements of TMPRSS2 activity in transfected cells. **A**. Western blot of TMPRSS2 protein expression in HEK-293T cells transfected with PLX304 vector or PLX304-TMPRSS2 vector is shown. TMPRSS2-containing V5 tag was assessed by anti-V5 antibody, and anti-GAPDH antibody was used as a loading control. **B**. Arbitrary fluorescence unit (AFU) measurements of control or TMPRSS-overexpressing cells incubated with BOC-QAR-AMC for 75 minutes at 37 °C are shown. Wells containing PBS and BOC-QAR-AMC were used as background fluorescence reads. **C**. Fluorescence of control or TMPRSS2-overexpressing cells was measured every 15 minutes for a total time of 180 minutes. The average proteolytic activity rate per minute **(D)** or hour **(E)** of control or TMPRSS2-overexpressing cells. The fluorescent signal was measured by the UV filter (excitation 365 nm and emission 410 nm). Data in B, D, and E represent the mean ± SD with interquartile ranges in B, D and E. GAPDH, glyceraldehyde-3-phosphate dehydrogenase; TMPRSS2, transmembrane serine protease 2.

### Identifying functional TMPRSS2 inhibitors

We tested the effect of protease inhibitors on TMPRSS2 activity. As a positive control, cells were treated with camostat mesylate, a drug that has been shown to inhibit TMPRSS2 (Hoffmann et al., 2020; Tim Hempel, 2020). As expected, camostat mesylate inhibited the proteolytic activity of TMPRSS2 with a calculated IC_50_ of 42 nM (Figure 2A). We then tested whether the secretory leukocyte protease inhibitor (SLPI) would inhibit TMPRSS2. However, none of the tested concentrations of SLPI inhibited TMPRSS2 proteolytic activity (Figure 2B). In contrast, A1AT inhibited TMPRSS2 proteolytic activity in a dose-dependent manner (IC_50_ of 357 nM; Figure 2C). A1AT did not demonstrate toxic effects in the tested concentrations as demonstrated by viability assays of HEK-293T cells (Supplementary Figure 2).

**Figure 2.**
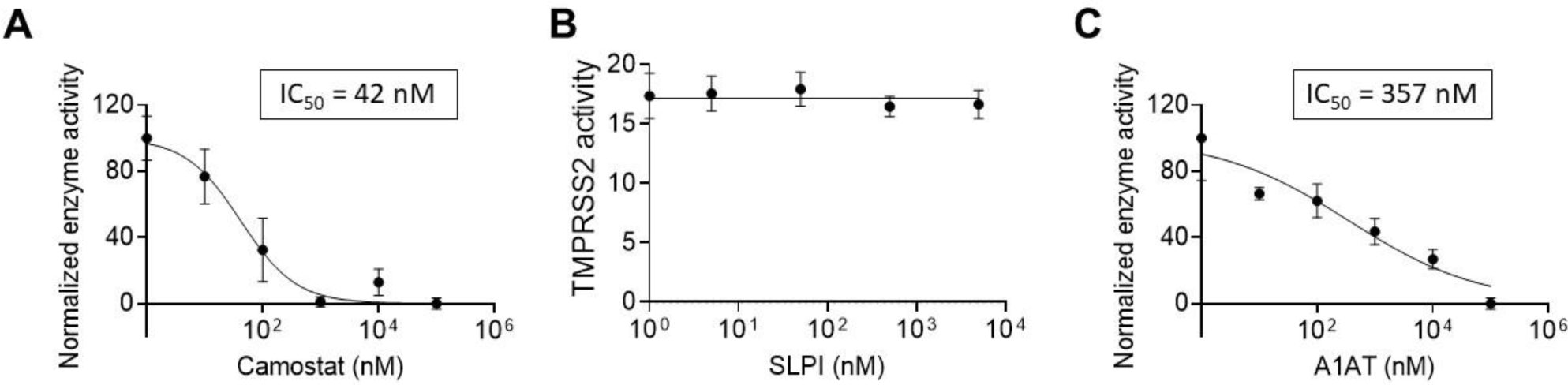
The effect of intracellular and extracellular inhibitors on TMPRSS2 activity. Fluorescence of TMPRSS2-overexpressing cells was measured every 15 minutes in the presence of the indicated concentrations of camostat mesylate (camostat) **(A)**, secretory leukocyte protease inhibitor (SLPI) **(B)**, or A1AT **(C)**. The results in A-C are presented as the means ± SE from at least 3 independent experiments performed in duplicate and/or triplicate. TMPRSS2, transmembrane serine protease 2.

### Modeling the extracellular TMPRSS2-A1AT Michaelis complex

We modelled the Michaelis complex between TMPRSS2 and A1AT to better understand the structural basis of TMPRSS2 inhibition prior to A1AT cleavage and covalent attachment (Figure 3A). Our results suggest that TMPRSS2 interacts with A1AT through its reactive center loop (RCL), driven by complementary electrostatic interactions at their surfaces (Figure 3B). Namely, LYS390 (TMPRSS2) forms a strong bifurcated salt bridge with GLU199/ASP202 (A1AT), while LYS340 and ASP260 form a second electrostatic contact, whose proximity may be limited by the presence of ASP338 at the surface of TMPRSS2. In the active site region, TMPRSS2 interacts with A1AT via an extensive hydrogen bond network (Figure 3C). Part of these interactions stabilize a short, antiparallel beta-sheet between GLY462 and ILE356-PRO357, similar to the one present in 1 OPH of the Research Collaboratory for Structural Bioinformatics (RCSB) Protein Data Bank (PDB) (Dementiev et al., 2003). At the entrance of the S1 pocket, GLN438 forms hydrogen bonds with PRO357 and SER359, fixing and orienting the A1AT backbone around the reactive peptide bond (MET358-SER359). As a result, MET358 is buried inside the S1 pocket, with its backbone carbonyl placed inside the oxyanion hole, forming hydrogen bonds with the GLY439 and SER441 (TMPRSS2) backbone amides. These predicted interactions, together with the overall orientation of the RCL and the catalytic triad, constitute a suitable environment for the cleavage and covalent binding of A1AT to TMPRSS2.

**Figure 3.**
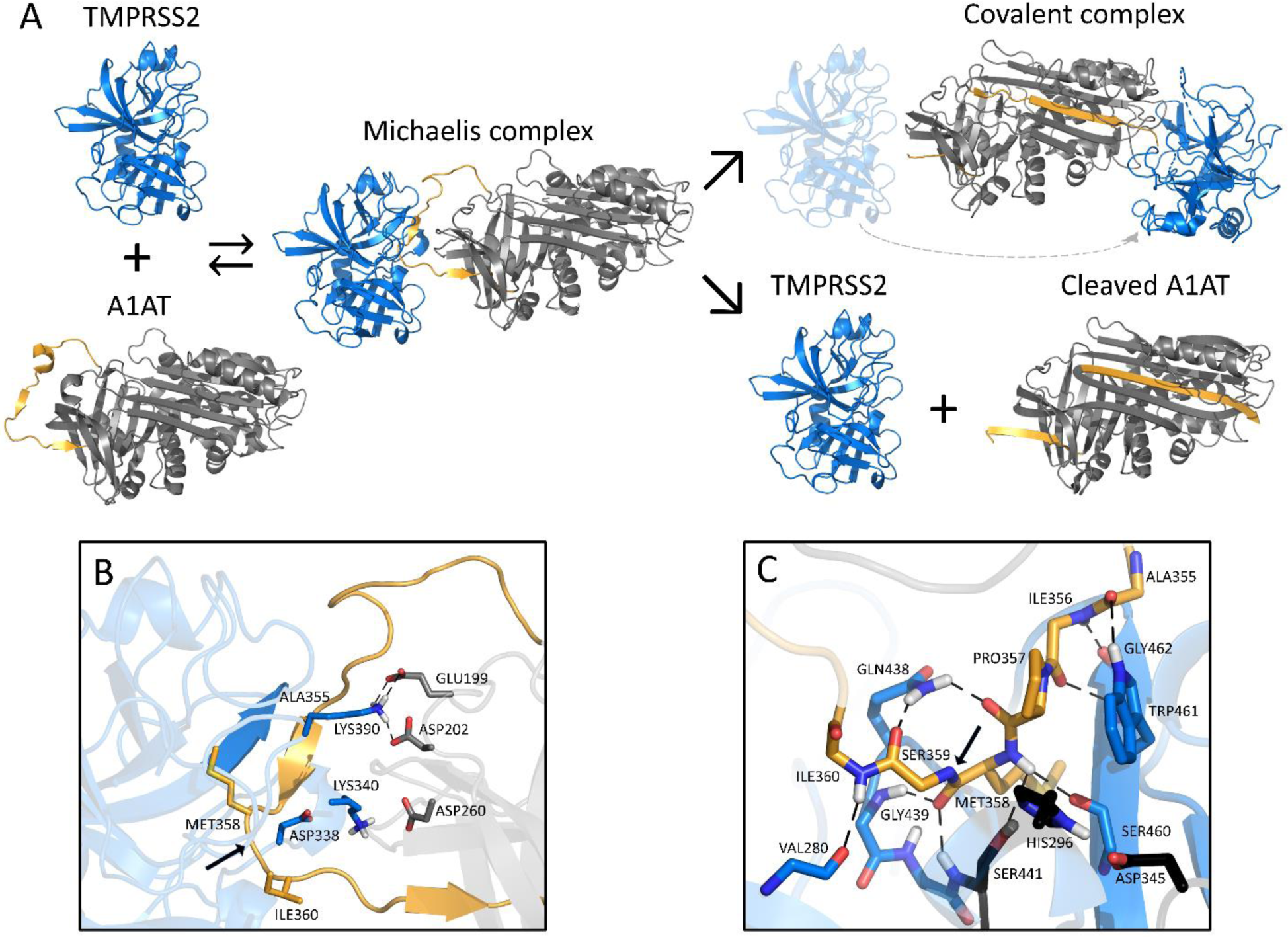
Schematic inhibition of TMPRSS2 by A1AT. **A**. General inhibitory mechanism of serpins applied to TMPRSS2 and A1AT are shown. The homology model of TMPRSS2 is shown as the blue cartoon and A1AT as the grey cartoon, with the reactive center loop highlighted in gold. **B**. Shown are the interactions at the interface of the Michaelis complex model, highlighting LYS340 and LYS390 of TMPRSS2 (blue) and GLU199, ASP202, and ASP260 of A1AT (grey). **C**. A close-up of the Michaelis complex at the active site region is shown. The catalytic triad residues HIS296, ASP345 and SER441 are depicted in black. Relevant residues are represented as sticks, hydrogen bonds are represented as dashed black lines, and the cleavage site is indicated by a black arrow. Note that there are hydrogen bonds at the oxyanion hole between GLY439/SER441 of TMPRSS2 and MET358 of A1AT. A1AT, alpha 1 antitrypsin; TMPRSS2, transmembrane serine protease 2.

### Analyzing the A1AT effect on TMPRSS2-mediated SARS-CoV-2 infectivity

We investigated whether A1AT can inhibit the entry of SARS-CoV-2 into cells that are commonly used for SARS-CoV-2 assays because of their TMPRSS2 expression (Hoffmann et al., 2020). Caco-2 cells were either left untreated or treated with either A1AT (10 µM) or camostat (10 µM); cells were then infected with SARS-CoV-2. Twenty hours later, quantifying the genomic SARS-CoV-2 from the intracellular RNA revealed a significant decrease in the viral load in cells that were treated with A1AT or camostat compared with untreated control cells (p < 0.0001 for both inhibitors; Figure 4). These data suggest that A1AT can limit the SARS-CoV-2 life cycle by modulating the host cells’ TMPRSS2 activity, similarly to camostat. Importantly, A1AT and camostat were nontoxic in Caco-2 cells in the tested concentrations (Supplementary Figure 3B).

**Figure 4.**
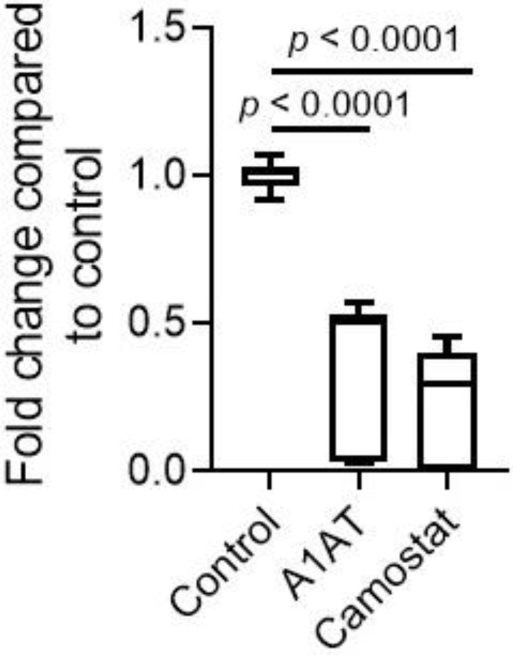
The effect of A1AT and camostat on SARS-CoV-2 genomic copies. Intracellular SARS-CoV-2 genomic copies were analyzed twenty hours after infection of Caco-2 cells in the presence of A1AT (10 µM), camostat (10 µM), or control media. Data represent fold change in CoV2 copy number compared to control media. Results are the mean ± SD of 3 independent experiments performed in duplicate and/or triplicate. A1AT, alpha 1 antitrypsin.

### Plasma A1AT levels in patients with COVID-19

A1AT is normally found at high concentrations in the blood and increases during acute phase responses or tissue injury (Guttman et al., 2015). We hypothesized that A1AT concentrations in plasma samples from patients with COVID-19 would correlate with disease severity as part of the anti–SARS-CoV-2 response. To test this hypothesis, we analyzed plasma A1AT levels in a cohort of patients who tested positive for COVID-19. These patients were divided into 3 groups according to disease severity at the time of emergency department disposition and the maximal severity within 30 days (1 mild – outpatient care, 2 moderate – need for hospitalization, 3 severe – need for intensive care unit admission; see methods section). A1AT levels were significantly different between the group of patients with mild disease and the group of patients with moderate disease. The mean concentration of A1AT was the highest in the group of patients with severe disease compared to the other groups (Figure 5A). A1AT concentrations positively correlated with maximal severity of disease (Figure 5B). Plasma A1AT concentrations correlated with plasma IL-6 (r = 0.65, p < 0.0001), IL-10 (r = 0.33, p = 0.002), and TNFα concentrations (r = 0.3; p = 0.002) but not plasma IL-8 concentrations (Figure 5C,D).

**Figure 5.**
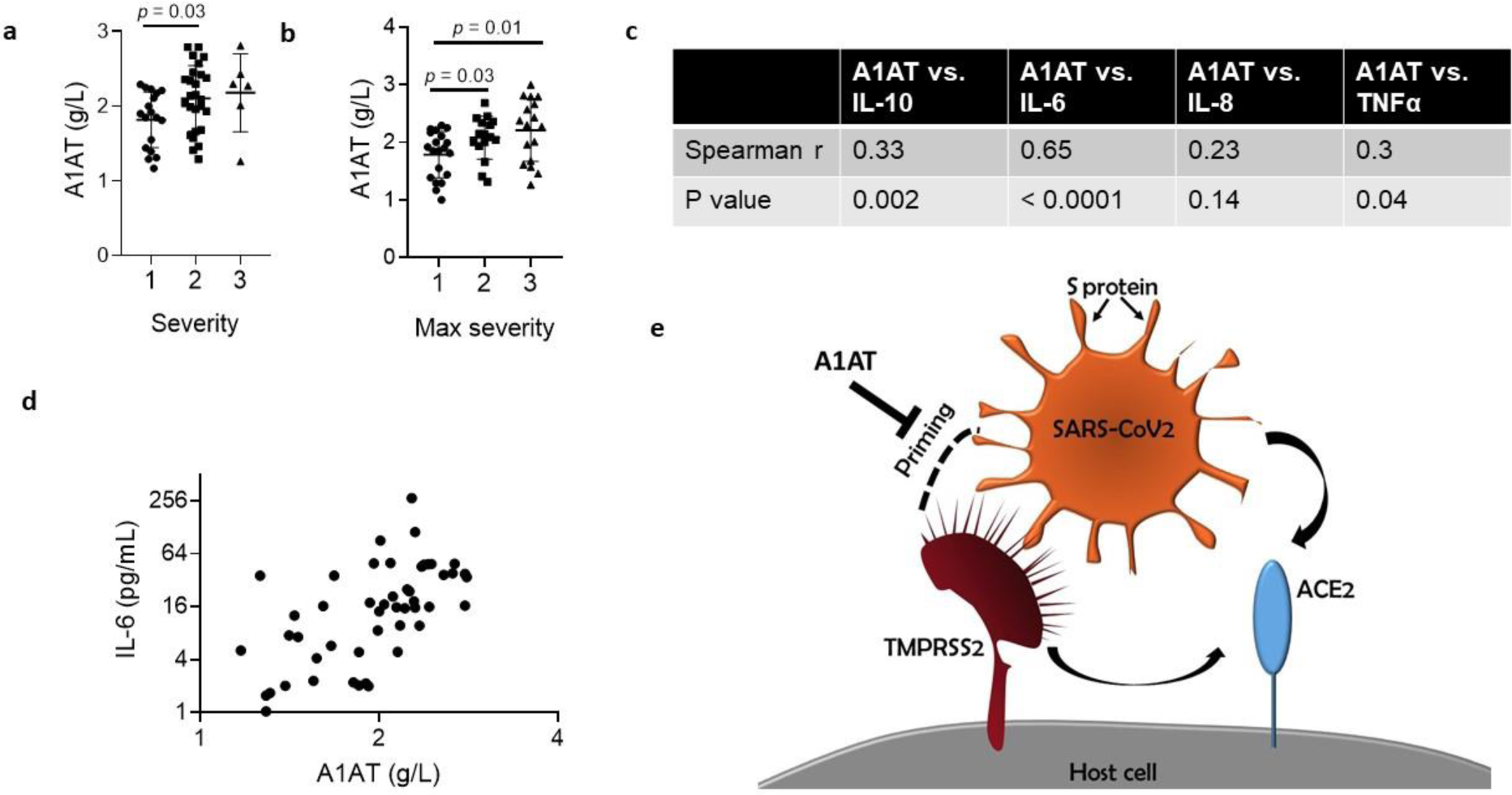
A1AT concentration in plasma samples of patients with COVID-19 and suggested role in SARS-CoV-2 cell entry. A1AT plasma concentrations in patients who were positive for COVID-19 are shown and stratified according to disease severity at the time of disposition from the Emergency Department **(a)** or the maximal severity within 30 days of index Emergency Department visit (Max severity) **(b)** (1 = Outpatient, 2 = Hospitalized, 3 = Intensive Care Unit or Death). In **(c)**, correlation between plasma A1AT concentrations and plasma IL-10, IL-6, IL-8, or TNFα concentrations. r and p values were calculated according to Spearman correlation. **d**. plasma concentration of A1AT and IL-6 in each patient with confirmed COVID-19 with markers representing individual patients. A1AT, alpha 1 antitrypsin; IL, interleukin. **e**. Model of SARS-CoV-2 entry mediated by extracellular proteolytic events. Extracellular proteases, such as TMPRSS2, process the S protein on the SARS-CoV-2 envelope in a process called priming. Priming of the S protein is necessary for binding between the S protein and the host receptor angiotensin-converting enzyme 2 (ACE2). Extracellular inhibitors, such as alpha 1 antitrypsin (A1AT), prevent the priming of the S protein and inhibit virus entry. In addition, inhibiting transmembrane serine protease 2 (TMPRSS2) prevents processing of ACE2, which decreases the infectivity of the coronavirus.

## Discussion

Herein, we developed a cell-based methodology that allows the quantification of TMPRSS2 activity. This methodology enables testing the effect of intracellular compounds and extracellular compounds, thus permitting differentiation between inhibition of intracellular and extracellular protease. Using this methodology, we revealed that A1AT, which is approved by the FDA for the treatment of A1AT deficiency, can efficiently inhibit TMPRSS2. Structural modeling of the A1AT-TMPRSS2 Michaelis complex revealed that A1AT is likely cleaved and covalently bound to TMPRSS2. The importance of A1AT in fighting coronavirus infection was supported by the finding that plasma A1AT levels correlated with COVID-19 severity and with plasma IL-6 levels. A1AT inhibited SARS-CoV-2 infection at a comparable level to camostat in Caco-2 cells, a cell type that is efficiently infected by SARS-CoV-2 (Chu et al., 2020). Therefore, our collective data suggest that A1AT treatment may benefit COVID-19 countermeasures by inhibiting extracellular-mediated S protein processing and virus entry.

Though the relative contribution of intracellular proteases and extracellular proteases to the S protein priming and cell entry of SARS-CoV-2 has yet to be determined, we provide evidence that extracellular protease activity is rate-limiting in the process of SARS-CoV-2 cell entry. Therefore, we suggest that extracellular protease inhibitor delivery may provide a good strategy for inhibiting SARS-CoV-2 entry and cell-to-cell transmission by modulating the exterior of the host cells. The disappointing results of hydroxychloroquine, which interfere with the activity of intracellular cathepsins in clinical trials and in vitro assays (Geleris et al., 2020; Kalligeros et al., 2020; Tianling Ou, 2020), are further consistent with our findings, which support the importance of extracellular proteases. Notably, neither A1AT nor camostat completely blocked SARS-CoV-2 entry. This could be explained by slow, unprocessed ACE2-mediated cell entry in the absence of TMPRSS2 and/or expression of other proteases, including intracellular proteases that may cleave the S protein.

Targeting the host extracellular proteases, such as TMPRSS2, has several advantages over targeting viral proteins. First, anti-virals can rapidly lose their effectiveness due to the high rate of mutations that occur in the viral genome, but targeting host proteins limits the risk of drug-resistant viruses due to the relatively low rate of mutations in the host genome. Notably, the obstacle in targeting human proteins is the potential risk of altering physiologic pathways. It was suggested that TMPRSS2 initiates a cascade of proteolytic activation events that regulate processing of proteins in seminal fluid and in the lung because TMPRSS2 regulates the sodium channel ENaC (Donaldson et al., 2002). Nevertheless, mice deficient in TMPRSS2 lack any obvious phenotypes, suggesting that other proteases may have redundant roles and may compensate for the loss of TMPRSS2 (Kim et al., 2006). Therefore, delivery of TMPRSS2 inhibitors during viral infections is likely a relatively safe strategy. Although the safety of TMPRSS2 inhibition has not been clinically proven yet, drugs with proteolytic inhibition activity towards TMPRSS2 (e.g., camostat mesylate and nafamostat mesylate) are currently being pursued for the treatment of COVID-19. Notably, unlike extracellular A1AT, camostat and nafamostat are cell permeable and therefore may possess undesired intracellular protease inhibition capacity. Second, inhibiting cell entry is an upstream intervention method that limits the overall viral burden and the spread to and replication within tissues, such as the salivary glands, that have important pathologic consequences involved in viral transmission to others. Third, inhibiting cell entry may prevent several downstream disease outcomes. For example, SARS-CoV-2–infected cells undergo cell death by pyroptosis, a process that is thought to induce complications, such as cytokine storm and intra-vascular thrombosis, and thereby result in severe disease outcomes. Moreover, cell death of key alveolar cells induces tissue damage that progresses into acute respiratory distress syndrome, a severe clinical phenotype of COVID-19. Therefore, inhibiting SARS-CoV-2 infection and cell entry by inhibiting the proteolytic priming of the coronavirus S protein is likely to decrease cell death and thereby reduce disease severity.

To our knowledge, we are the first to demonstrate that A1AT inhibits TMPRSS2, which is an extracellular protease with a key role in the entry of SARS-CoV-2, SARS-CoV, MERS-CoV, and influenza viruses (Gierer et al., 2013; Harbig et al., 2020; Hatesuer et al., 2013; Hoffmann et al., 2020; Sakai et al., 2014; Shirato et al., 2013). A1AT belongs to the super family of serine protease inhibitors (SERPIN) that irreversibly inhibit serine and cysteine proteases. Proteases interact with SERPINs as depicted in Figure 3A, forming a Michaelis complex in which the reactive center loop (RCL) binds the protease active site (modelled for TMPRSS2 and A1AT in this work). Cleavage of the RCL results in the formation of a transient, covalent complex that can either undergo dissociation (lower right pathway in Figure 3A; cleaved A1AT represented by PDB 7API (Engh et al., 1989)) or translocation and irreversible inhibition (Bao et al., 2018) (upper right pathway in Figure 3A; represented by PDB 2D26 (Dementiev et al., 2006) in the absence of a TMPRSS2-A1AT– specific model). One of the fragments of the cleaved RCL is inserted into the central beta-sheet of A1AT in both pathways, whereas the other fragment (36-aa) is released into the solvent. Notably, the 36-aa fragment is produced as a result of proteolytic cleavage by several serine proteases and possesses physiologic functions (Blaurock et al., 2016; Potere et al., 2019; Risor et al., 2017).

A1AT concentration in the blood can be increased by 6 fold as part of the acute phase of inflammation or tissue injury (Guttman et al., 2015). A1AT is known to inhibit neutrophil elastase, proteinase 3, and cathepsin G. Neutrophil elastase cleaves several structural proteins in the lungs, processes several innate immune mediators, and has been shown to be involved in the pathogenicity of SARS-CoV-2 substrains by cleaving the S1-S2 junction of the S protein (Chandrika Bhattacharyya, 2020). In addition, A1AT promotes clearance of apoptotic cells (Serban et al., 2017). Increased neutrophil levels have been found in patients with COVID-19 with severe disease compared to those with mild disease and healthy controls (Henry et al., 2020; Qin et al., 2020; Wang et al., 2020). Notably, truncated forms of A1AT were significantly increased in the serum of patients with SARS compared to control patients, and the truncated A1AT levels correlated with disease severity (Ren et al., 2004). These findings suggest that A1AT may be a part of a natural protective mechanism to fight coronavirus infection and acute lung disease. Indeed, we demonstrated that plasma A1AT levels were associated with COVID-19 severity and with IL-6, a cytokine that has been implicated in COVID-19 pathology and as a biomarker for disease severity (Chen et al., 2020; Henry et al., 2020; McElvaney et al., 2020; Ulhaq and Soraya, 2020; Zhou et al., 2020). Notably, A1AT was shown to inhibit the infection of H3N2 influenza A and influenza B viruses in a murine model (Harbig et al., 2020), even though these viruses do not require TMPRSS2 priming, underscoring that A1AT can mediate anti-viral effects via multiple mechanisms.

We suggest that treatment with extracellular protease inhibitors either alone or in combination with other anti–COVID-19 agents may be a useful antiviral strategy to fight COVID-19. These protease inhibitors have the potential to prevent SARS-CoV-2 entry to host cells by inhibiting S protein priming by TMPRSS2 and other extracellular proteases and binding of the virus to ACE2 (Figure 5E). A1AT may be particularly effective as it has dual capacity, inhibiting TMPRSS2 (and hence viral uptake and subsequent replication) and possessing anti-inflammatory activity(Janciauskiene and Welte, 2016). We suggest that using these inhibitors may be therapeutic in conditions in which TMPRSS2 function is pathogenic, such as in several types of coronavirus and influenza infections. The ready availability of and safety profile A1AT calls attention to its potential clinical use for the COVID-19 pandemic.

## Materials and Methods

### Materials

Secretory leukocyte peptidase inhibitor (SLPI) and Boc-Gln-Ala-Arg-7-Amino-4-methylcoumarin (BOC-QAR-AMC) were obtained from R&D systems. Camostat mesylate was obtained from Sigma Aldrich, and A1AT (CSL Behring, Zemaira and Grifols, Prolastin-C) was a kind gift of Mark Brantly (University of Florida, Gainesville, FL). Recombinant TMPRSS2 was obtained from Abnova.

### TMPRSS2 overexpression

A PLX304 plasmid–containing human *TMPRSS2* open reading frame from the ORFeome Collaboration (Dana-Farber Cancer Institute, Broad Institute of Harvard and Massachusetts Institute of Technology [<u>HsCD00435929</u>]) was obtained from DNASU Plasmid Repository, and a control PLX304 vector was obtained from Addgene (Watertown, MA, USA).

### HEK-293T cell culture and transfection

HEK-293T cells were grown in Dulbecco’s modified eagle media (DMEM) supplemented with 10% fetal bovine serum (FBS) and seeded in a black, 96-well plate (75,000 cells/well). The following day, cells were transfected overnight with either a control plasmid (PLX) or TMPRSS2 (PLX-TMPRSS2) via TransIT LT-1 transfection reagent (Mirus Bio) in 100 µL of OptiMEM per well. The media was replaced the day following the overnight transfection.

### TMPRSS2 activity assay

Twenty-four hours after transfection, the media was replaced with 80 µL of phosphate-buffered saline (PBS). Inhibitors or PBS alone were added to the wells in the indicated concentrations and incubated at 25°C for 15 minutes. The fluorogenic substrate BOC-QAR-AMC (R&D Biosystems) was then added to each well to a final concentration of 100 µM. Fluorescence (excitation 365 nm, emission 410 nm) was immediately measured every 15 minutes at 37°C using a GloMax plate reader (Promega).

### Gel electrophoresis and western blot

Protein lysates of HEK-293T cells were extracted with RIPA buffer (PIERCE) and protease inhibitor cocktail (Roche). Loading buffer (Life Technologies) was added, and samples were heated to 95°C for 5 minutes and subjected to electrophoresis in 12% NuPAGE Bis-Tris gels (Life Technologies). Gels were transferred to nitrocellulose membranes (Life Technologies) and probed with the primary antibodies rabbit anti-V5 (Bethyl Laboratories), mouse anti-TMPRSS2 (Santa Cruz), and rabbit anti–human GAPDH (ABCAM) and subsequently with the secondary antibody IRDye 800RD goat anti-rabbit and IRDye 680RD goat anti-mouse (LI-COR Biosciences). Membranes were visualized and analyzed using the Odyssey CLx system (LI-COR Biosciences). For TMPRSS2 quantity estimations, recombinant TMPRSS2 was subjected to gel electrophoresis with protein lysates from TMPRSS2-overexpressing cells.

### Cell viability assay

HEK-293T cells were grown for 48 hours in 96-well plates. Then cells were treated with the indicated concentrations of A1AT for an additional 3.25 hours. Cell viability was assessed by Alamar blue cell viability reagent (ThermoFisher) according to the manufacturer’s protocol.

Caco-2 cells were plated in 96-well plates. Cells were treated with A1AT or camostat at the indicated concentrations for 18 hours. Cell viability was estimated by CellTiter-Glo Cell Viability Assay from Promega according to the manufacturer’s protocol.

### Quantifying SARS-CoV-2 genomic copies

Caco-2 cells were plated in 12-well plates and pretreated for 1 hour prior SARS-CoV-2 infection with DMEM containing A1AT or camostat in the indicated concentrations or DMEM alone (media-only control). Cells were incubated with SARS-CoV-2 (through BEI Resources, NIAID, NIH: SARS-Related Coronavirus 2 Isolate USA-WA1/2020, NR-52281) at a multiplicity of infection of 0.1 and with the indicated drugs for 1 hour. Then, cells were washed with DMEM and incubated with DMEM containing A1AT or camostat or DMEM alone. Twenty hours post infection, RNA was extracted from the cells, and viral copy number was quantified by real-time, quantitative PCR.

### mRNA extraction and real-time, quantitative PCR

Cells were lysed with TriZol LS, and total RNA was isolated from cells with the Direct-zol RNA mini prep kit (Zymo research) according to the manufacturer’s protocol. For RNA sequencing experiments, RNA was treated with the On-Column DNase Digestion kit (Qiagen) according to the supplied protocol. The amount of the intracellular RNA was estimated by real-time quantitative PCR performed with the RNA UltraSense™ One-Step Quantitative RT-PCR System (Applied Biosystems) by using the following primer sets: *CDC N1* (forward 5′-GAC CCC AAA ATC AGC GAA AT, reverse 5’-TCT GGT TAC TGC CAG TTG AAT CTG, probe 5’-FAM-ACC CCG CAT TAC GTT TGG TGG ACC -BHQ1), *18S rRNA* (TaqMan Gene expression assay from ThermoFisher, Hs99999901_s1), and *GusB* (TaqMan Gene expression assay from ThermoFisher, Hs99999908_m1).

### Template-based molecular docking

The Michaelis complex of TMPRSS2 and A1AT was modelled using HADDOCK v2.4 (Dominguez et al., 2003), a versatile docking engine guided by structural data from experiments and/or computational models. A complex between S195A bovine cationic trypsin and the Pittsburgh variant of A1AT (PDB 1OPH (Dementiev et al., 2003)) was taken as a template to drive the docking process, as well as to obtain the structure of A1AT (chain A, reverted to its wild-type form). The structure of TMPRSS2 was obtained from reference (Rensi et al., 2020) (homology model based on PDB 3W94). PS-HomPPI v2.0 (Xue et al., 2011) was used for mapping the interface residues of PDB 1OPH onto the query structures and calculating their CA-CA distances. A subset of these distances within different cutoffs (7, 9, 10, 11, and 15 Å) was then used as unambiguous restraints to drive the docking between TMPRSS2 and A1AT. The number of structures generated in each of the docking phases, i.e., rigid-body, semi-flexible, and water refinement, was kept at 1000, 200, and 200, respectively. The top 4 models generated using each of the 5 cutoffs were analyzed, and the final model was selected on the basis of visual inspection and energetic terms.

### COVID-19 plasma analysis

Adults who presented to the University of Cincinnati Medical Center (UCMC) Emergency Department (ED) with suspected COVID-19 and had a clinically indicated blood draw were prospectively enrolled via an institutional review board–approved waiver of informed consent. Following collection, samples were centrifuged at 2000 g for 15 min at 4°C and frozen at −80°C until analysis. Inclusion in this analysis was dependent on a positive reverse transcription polymerase chain reaction (RT-PCR) test for COVID-19 via a standard-of-care nasopharyngeal swab. A total of 49 patients were included. The median age was 46.5 years (IQR: 37–66). The plasma concentration of A1AT was measured on a Behring Nephelometer II System (BN II, Siemens Medical Solutions USA, Inc., Malvern, PA, USA). The plasma concentrations of IL-6, IL-8, IL-10, and TNFα were quantified using the Meso Scale Discovery (MSD) U-Plex assay (Rockville, Maryland, USA). Patients who were positive for COVID-19 were stratified into subgroups on the basis of disease severity at ED disposition and maximal disease severity within 30 days of the index ED visit. Mild COVID-19 was defined as illness requiring only outpatient care (level 1), moderate COVID-19 was defined as illness requiring hospitalization (level 2), and severe COVID-19 was defined as illness requiring Intensive Care Unit admission and/or mechanical ventilation and/or resulted in the death of the patient. Comparing plasma concentrations of A1AT with those of IL-10, IL-6, IL-8, and TNFα were performed using the Spearman’s test.

### Statistics

IC_50_ values were calculated by nonlinear regression by dose-response inhibition with variable slopes. Statistical significance was determined using a t test (unpaired, two-tailed). Statistical analyses were performed using GraphPad Prism (GraphPad Software Incorporated).

## Author contributions

N.P.A. designed and performed experiments and data analysis and wrote the paper, A.M.K. performed experiments, V.C. and I.V.A. performed experiments with SARS-CoV-2. K.E. and L.R. assisted in data generation and analysis of template-based molecular docking. B.M.H., J.L.B., and S.W.B. conducted patient sample collection and plasma tests of patients with COVID-19. F.N. supervised the template-based molecular docking. K.K-H. supervised the SARS-CoV-2 experiments. M.E.R. supervised the study.

## Acknowledgments

We thank M. Brantly (Alpha-1 Foundation) for the gift of A1AT and B.J. Aronow, J.D. Molkentin, and S.N. Waggoner (Cincinnati Children’s Hospital) for insightful discussion and advice. We also thank S. Hottinger (Cincinnati Children’s Hospital) for editorial assistance. The following reagent was deposited by the Centers for Disease Control and Prevention and obtained through BEI Resources, NIAID, NIH: SARS-Related Coronavirus 2 Isolate USA-WA1/2020, NR-52281.

## Supplementary Information

**Supplementary Figure 1.**
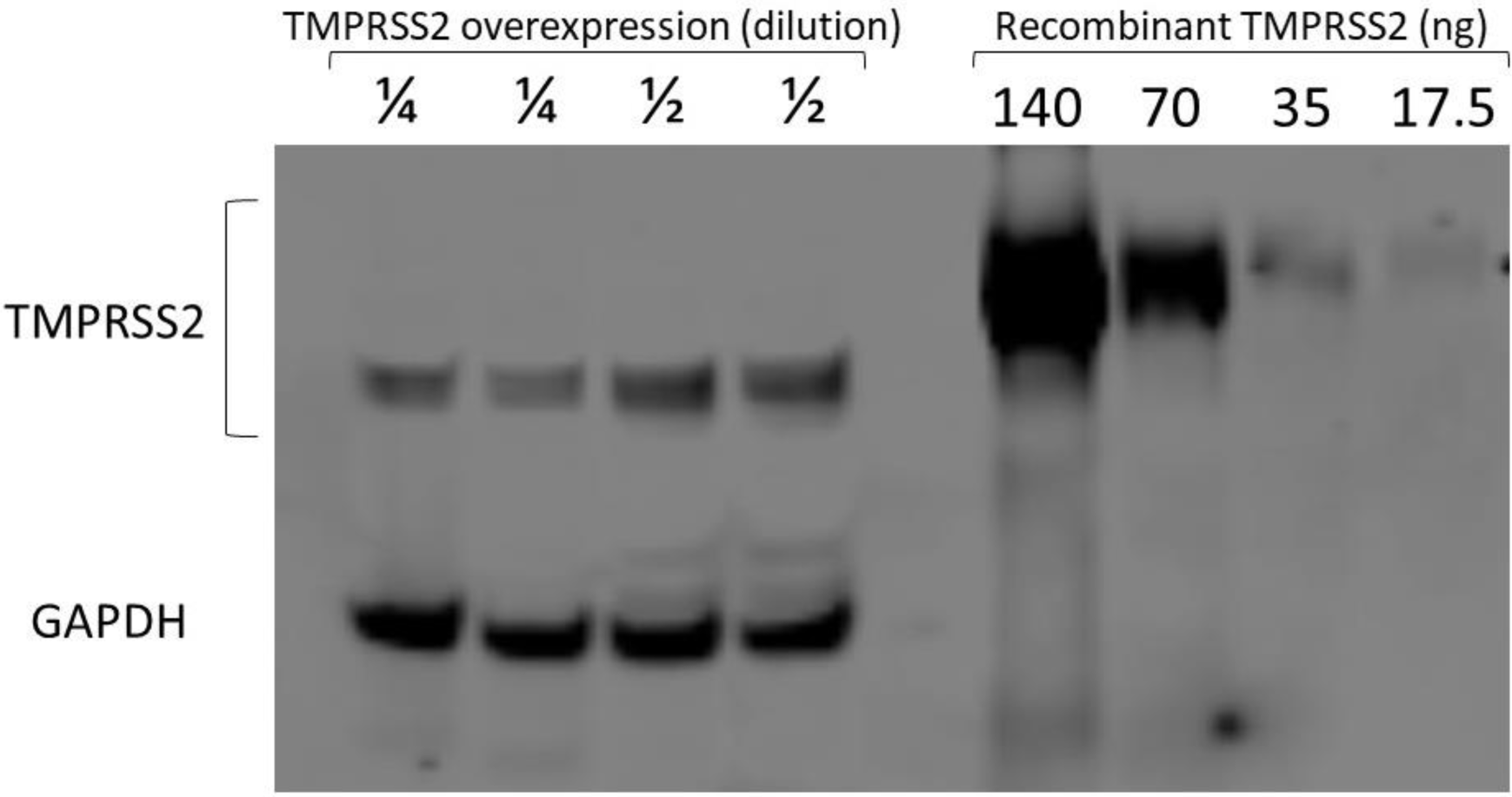
TMPRSS2 protein quantity calculation. Western blot analysis of protein lysates from TMPRSS2-overexpressing cells (transfected HEK-293T) that were diluted ¼ or ½ and of recombinant TMPRSS2 as indicated. GAPDH, glyceraldehyde-3-phosphate dehydrogenase; TMPRSS2, transmembrane serine protease 2.

**Supplementary Figure 2.**
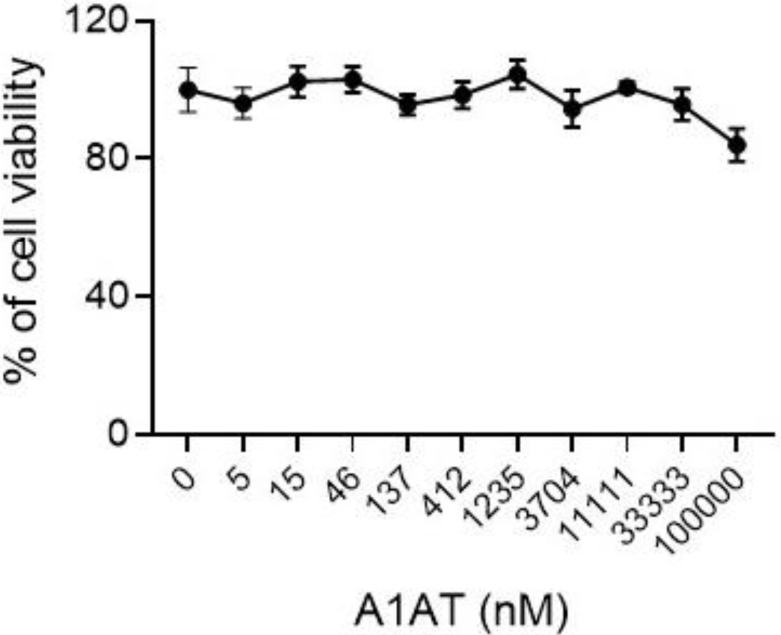
Viability assay. HEK-293T cells were assessed according to their viability after 18 hours at the indicated concentrations of A1AT. Cell viability was calculated as the percent of viability compared to untreated cells. Results are presented as the mean ± SE of 2 independent experiments performed in 6 replicates. A1AT, alpha 1 antitrypsin.

**Supplementary Figure 3.**
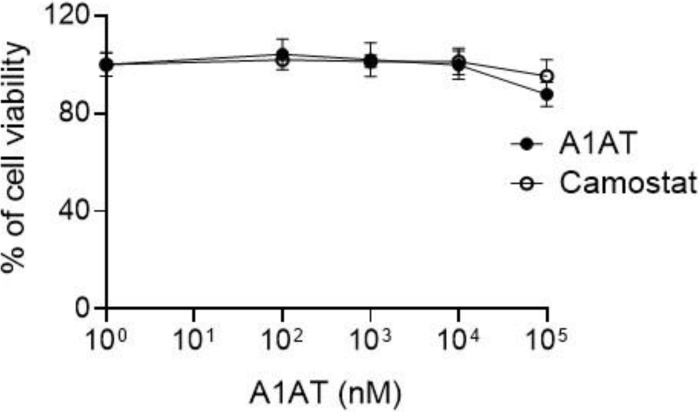
Viability assay of A1AT and camostat on SARS-CoV-2 genomic copies. Caco-2 cells were assessed according to their viability at the indicated concentrations of A1AT. Cell viability was calculated as the percent of viability compared to untreated cells. Results are the mean ± SE. A1AT, alpha 1 antitrypsin; camostat, camostat methylate.

